# Investigating Data Size, Sequence Diversity, and Model Complexity in MPRA-based Sequence-to-Function Prediction

**DOI:** 10.1101/2025.03.11.642630

**Authors:** Yilun Sheng, Xinming Tu, Sara Mostafavi

## Abstract

We created the MPRA Dataset Collection (MDC), a curated resource of MPRA data from 12 studies comprising over 150 million labeled DNA subsequences. These datasets include both random and natural genomic sequences paired with diverse functional outputs such as gene expression and splicing efficiency. Using this collection, we build sequence-to-function (S2F) predictive models of regulatory elements and analyze these models to uncover insights into the relationship between training data requirements, experimental design, and model generalizability. By offering a high-quality, machine learning-ready repository, MDC accelerates the development of robust computational tools for deciphering the mechanisms of gene regulation.

## Main

Massively Parallel Reporter Assays (MPRAs) generate datasets that are important for training and making inferences from sequence-to-function (S2F) models of gene regulation^1,2^. By including synthetic sequences outside the natural DNA distribution, they provide a rigorous test of model generalizability^2–4^. Moreover, the large-scale data generated by MPRA experiments—on the order of millions of labeled DNA subsequences—matches the data demands of modern deep learning methods^5^, making MPRAs an ideal platform for both training and validating predictive models of gene regulation^2,6^.

Despite their advantages, large-scale and cross-study analysis of MPRA datasets are hindered by the lack of standardized, machine learning (ML)-ready datasets. While previous efforts have aimed to aggregate and standardize MPRA datasets to facilitate broader analyses and applications^7–9^, these efforts often lack direct integration with machine learning workflows or do not encompass synthetic and out-of-distribution sequences critical for probing regulatory functions beyond natural constraints.

Here, we developed the MPRA Dataset Collection (MDC), a curated resource tailored to support ML applications in functional genomics. MDC was put together by a systematic literature search and curation, resulting in datasets from 12 peer-reviewed studies (**Table 1, Figure 1a**), together encompassing >150M genomic DNA sequence matched with their functional output (see Method). Specifically, the MDC encompasses short genomic DNA (50bp-250bp) that are paired with “labels”, which we categorized into four groups based on the type of regulatory component assayed. The majority of these include random sequences tested for their ability to generate gene expression at promoters (n = 123,097,484)^4,5,10,11^. MDC also includes random and natural exonic sequences, tested for their splicing efficiency (n = 6,905,101)^3,12,13^, as well as random and natural 5’ UTR sequences tested for their translational efficiency (n = 3,155,072)^14,15^. As well, this collection includes native candidate regulatory elements (candidate enhancers) tested for their ability to drive gene expression in cell lines (n = 360,779)^10,16–18^. To enable wide utility of this resource, we developed an easy-to-use Python package for transforming the raw-data into ML-ready format, which also integrates numerical libraries including Pandas and PyTorch to support S2F prediction workflows (see Methods, **Figure 1b**).

**Figure 1.**
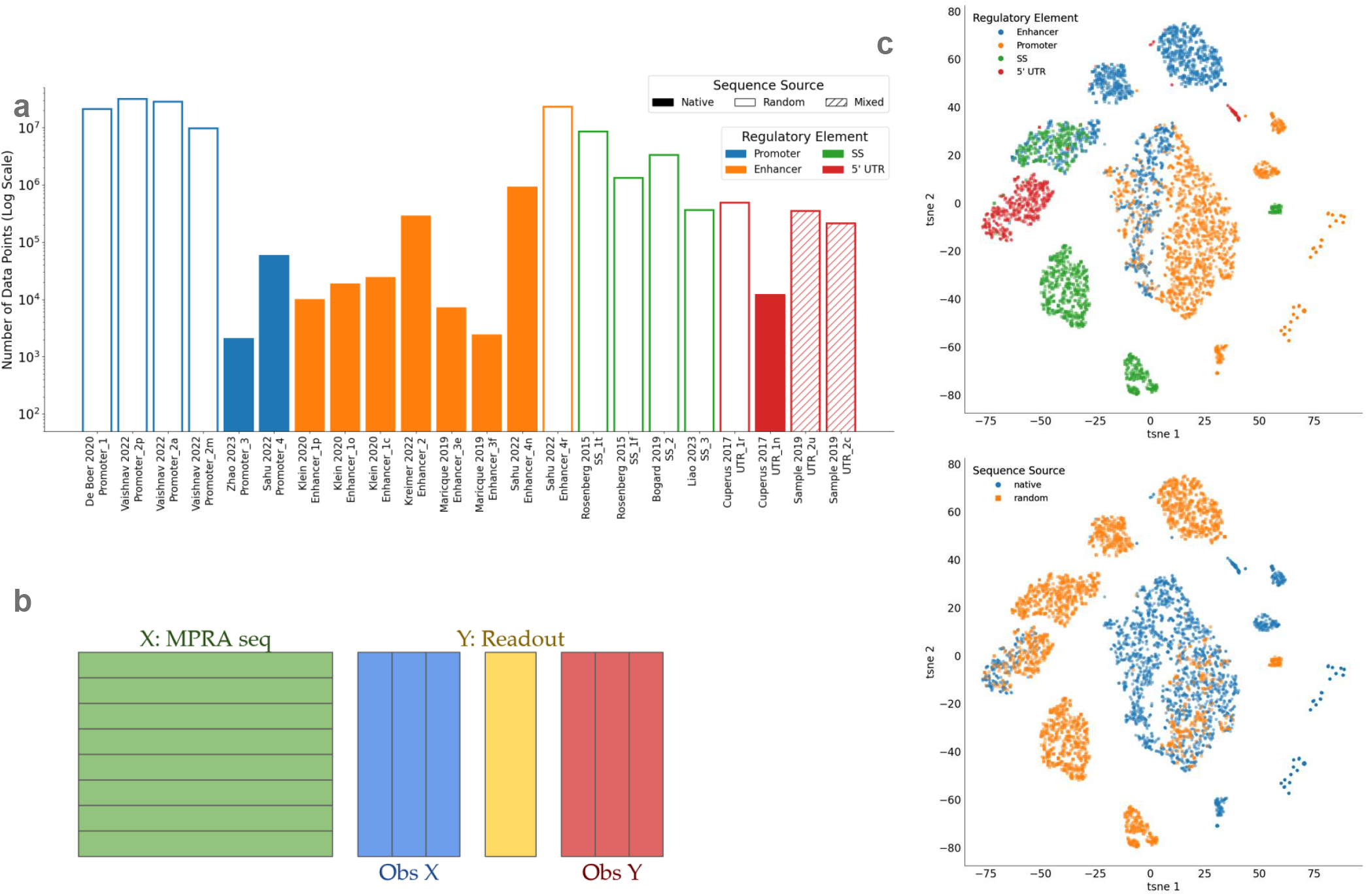
Representation of the MPRA Data Collection. a) Bar chart showing the number of DNA sequences captured by each study including in MDC. b) MDC software package consisted of unstructured meta-information and structured dataset. Each row is associated with one unique test sequence X (green) and one or more readout values Y (yellow). Observations on either X or Y were stored in extra columns (blue and red). c) Embedding from application of HyenaDNA to sequences in the MDC. Each dot represents one genomic DNA sequence. HyenaDNA was applied to each sequence, and the embedding space representation was extracted. This representation was visualized in 2D using t-SNE^22^. Top: data points are colored by the type of regulatory element they were tested for in MDC collection (SE: splicing efficiency). Bottom: data points are colored by their origin, either random or native.

To assess the diversity of genomic DNA represented by MDC, we applied existing DNA language models, including HyenaDNA^19^, to visualize the functional representation of this collection. As shown in **Figure 1c**, the embedding space captured by HyenaDNA indicates that MDC’s sequences encompass a broad range of functional elements, spanning both random and native sequence space. As nicely captured by HyenaDNA representations, random DNA broadens the sequence representation space and occupies a distinct region compared to native DNA.

We next explored two key open questions in functional genomics with S2F models^1,6^: (1) how training data size influences model generalization and (2) whether random DNA sequences provide similar or greater data efficiency compared to native genomic DNA.

To address the first question, we drew on the concept of *scaling laws* from deep learning, examining how model performance changes as we vary the amount of training data for each dataset. This approach helps reveal the intrinsic difficulty of each dataset and type for learning sequence-to-function relationships. Concretely, for each dataset, we kept a fixed test set (∼20% of the full data), and varied the size of the training set by sub-sampling. We then trained a CNN-based sequence-to-function (S2F) model (fixed architecture) on the sub-sampled training sets, enabling us to assess how training sample size influences performance (see Methods).

**Figure 2a** shows the result for representative datasets (here we performed analysis only on those datasets with sufficient reproducibility; **Figure S1a, Table 3**). As expected, increasing training size improves model performance, and at certain training sizes the model’s performance starts to plateau. The interesting component here is the speed at which the model reaches its near-optimal performance (if at all). For example, **Figure 2a** shows that predicting promoter expression and splicing outcomes requires less training data compared to enhancer expression. These differences highlight the unique properties of each dataset, and can be attributed to two factors: noise in the experiment, and the biological complexity of the underlying regulatory code.

**Figure 2.**
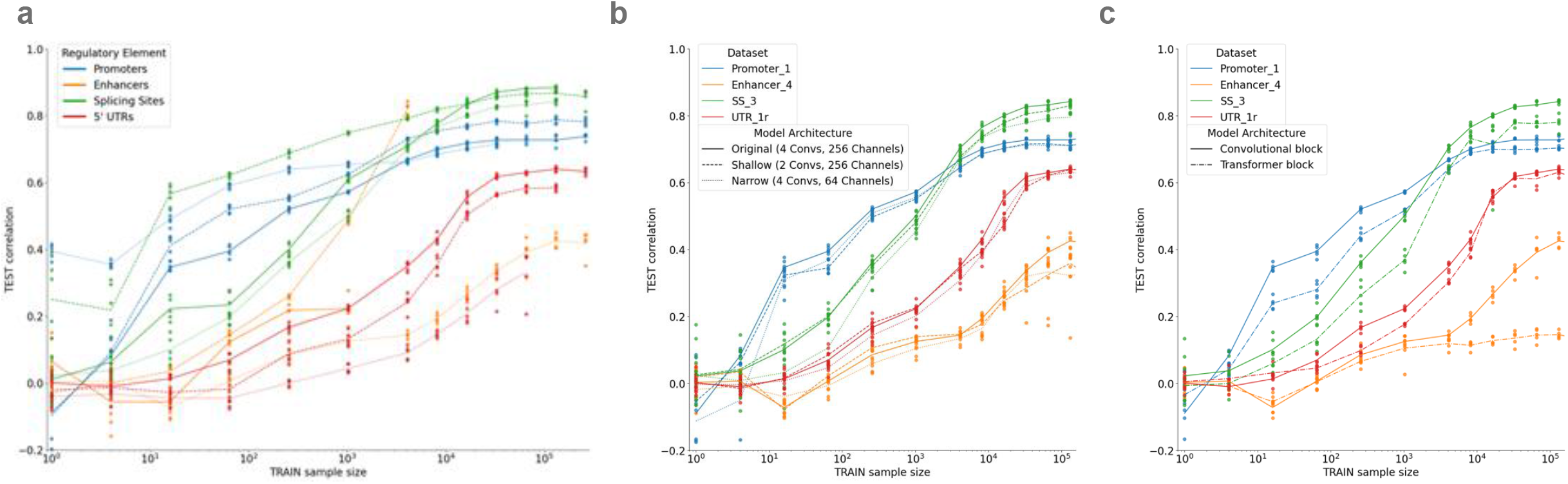
The relationship between training size, task difficulty, and prediction performance. a) The x-axis represents the number of training samples (logarithmic scale), the y-axis shows the test set correlation between observed and predictions. Results are shown for the best model selected based on the validation set. Each curve corresponded to a specific dataset, with line colors representing the type of regulatory element studied (e.g., promoters, enhancers, splicing sites, or 5’ UTRs). b) Different line styles represent variations in model architecture: dashed lines correspond to variations on convolutional blocks from 4 to 2 (shallow model); dotted indicate variations in the number of channels (kernels) from 256 to 64 (narrow model). c) Different line styles represent variations in model architecture: solid lines indicate model with Convolutional blocks, and dashed lines indicate model with Transformer blocks. See **Table 3** for datasets showcased here.

To systematically compare datasets, the above analysis uses a fixed model architecture. This choice, however, can impact the nature of the scaling law curves (if for example, a model architecture is a better fit for one dataset than another). Thus, we examined the effects of varying model architectures on scaling law curves. In particular, we altered the model’s complexity in two ways: 1) reducing model depth (number of convolutional layers) and width (number of channels in each layer) of CNNs (**Figure 2b, S2b-c**); 2) including transformer-blocks (**Figure 2c**). We made some notable observations: generally, increasing model depth has a larger impact on performance, compared to model’s width; transformer-blocks are generally not that performant when modeling short genomic DNA and tend to result in underperformance with limited dataset size. In summary, we observed that model architecture changes introduced minor shifts in the scaling law curves, and their effects were marginal compared to the differences observed between datasets.

Next, to gain deeper insights into how S2F models scale with different types of input data, we examined the performance of models trained on both natural (native) and synthetic (random) 5’UTR sequences using the dataset generated by Cuperus et al. 2017^14^. This unique dataset contains native 5’UTRs (n = 11,962) and randomized 50-nucleotide segments (n = 489,242). **Figure 3a** shows a four-way comparison in which each model is trained on either native or random sequences and then evaluated on native or random sequences. Somewhat counterintuitively, models trained on native sequences perform well with fewer training examples when predicting on native data. In contrast, while models trained on random sequences are initially less data-efficient, they eventually surpass the native-trained models with larger training sets. One explanation is that random DNA contains more “non-functional” signals, making it harder to learn from smaller datasets but potentially providing a broader range of patterns that ultimately improve generalization.

**Figure 3.**
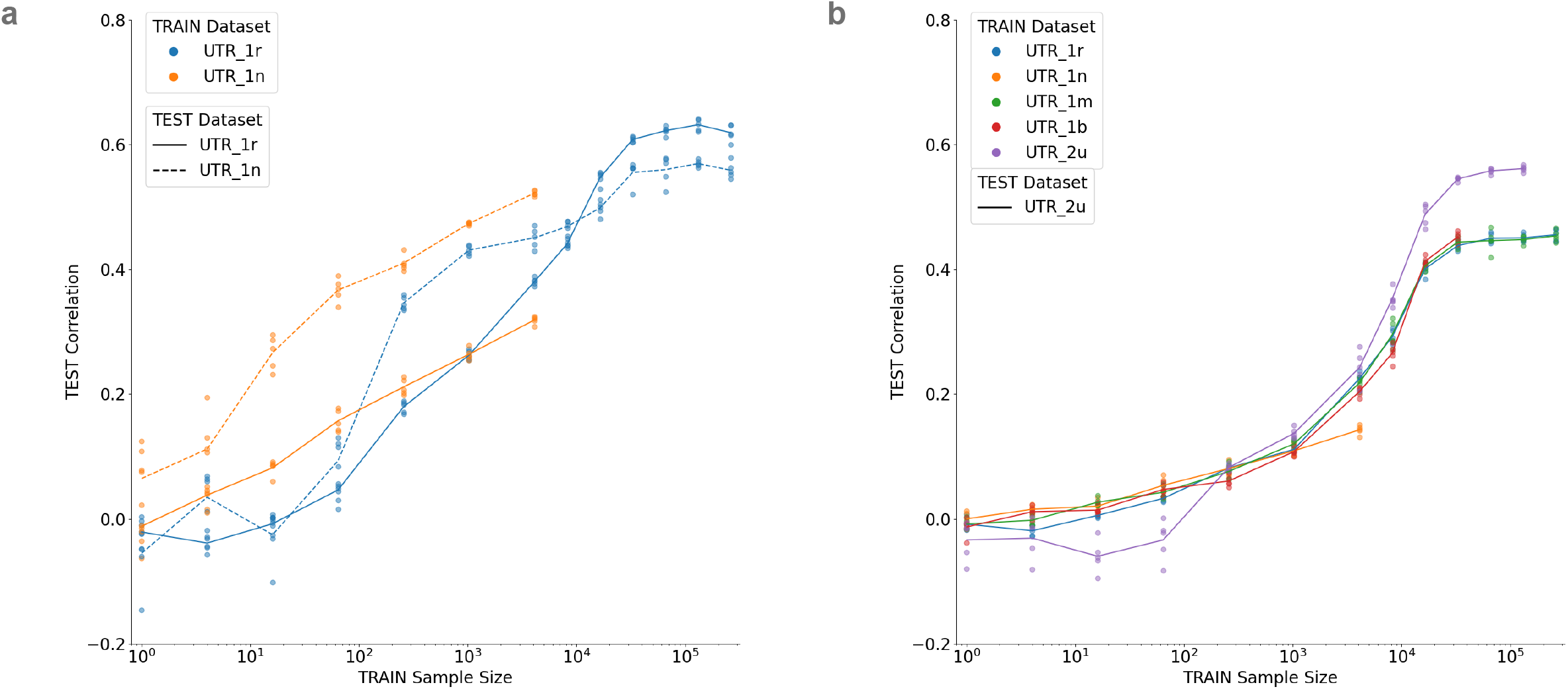
Data efficiency for learning S2F models using native vs. random DNA sequences. a) This plot showed the test set correlation as a function of training data size. Models were trained UTR-tested sequences from Cuperus et al., 2017^14^. Four models were trained and tested: models were trained on random DNA (UTR_1r, blue) or native DNA (UTR_1n, orange) and tested either on random DNA (solid lines) or native DNA (dashed line). For instance, the blue dashed line shows the performance of models trained with varying sample sizes of random DNA to predict UTR expression, and tested on the random DNA. b) Orange (UTR_1n), blue (UTR_1r), green (UTR_1m) and red (UTR_1b) shows the test set correlation (y-axis) when yeast-trained models are evaluated on the human UTR dataset^15^. The training size on the x-axis refers to the training of “yeast models”. The four yeast models include: a model trained only on random DNA (blue), a model trained only on native DNA (orange), and two hybrid models using both random sequence and native yeast sequences. The green is a hybrid model trained on training data that contains 98% random and 2% native native sequences. The red is a hybrid trained model with 80% random and 20% native sequences. The purple curve shows the performance of a model trained and tested on human UTR data, to set the expected range of performance. See **Table 4** for complete information on datasets used in this experiment.

Next, we took advantage of the availability of human UTR MPRA dataset (n = 351,575) by Sample et al. 2019^15^ in our MDC, and further tested the “out-of-distribution” generalizability of models trained on native or random DNA from yeast. **Figure 3b** shows the prediction performance of S2F models trained on yeast UTRs with varying training sample sizes. To establish an upper bar on expected performance, we also performed similar scaling law experiments using a human UTR dataset for model training and testing. Models trained on both random and native yeast UTRs (blue and orange curves) show competitive performance, reaching the upper bar in the low-data regime. However, “yeast-trained” models plateau below the level achievable with human data (purple curve), reflecting inherent differences in experimental settings. Finally, we trained models on a mixture of random and native yeast sequences (green and red curves) to test whether a larger amount of yeast training data could close the cross-species gap; the results suggest that beyond a certain point, increasing yeast data is unlikely to further improve performance.

In summary, the MDC addresses an important need in regulatory genomics: a standardized, ML-ready repository of MPRA data that covers diverse sequence types and experimental conditions. MDC simplifies access to large-scale MPRA datasets through a uniform interface, enabling researchers to systematically compare how different types of regulatory elements inform sequence-to-function prediction. Enabled by MDC, our analyses underscore several important takeaways for S2F modeling. First, we showed quantitatively that larger datasets generally improve model performance, but their impact varies by regulatory element. This suggests that some elements, like promoters, may possess more straightforward sequence features than others, such as enhancers. Second, we observed that while deeper CNNs and transformer-based models can bring incremental gains, the most pronounced performance differences arise from the biological and experimental properties of each dataset. Third, we observed that native sequences are more data-efficient when one has a limited number of training data points, presumably because they capture biologically relevant patterns. However, synthetic sequences can provide broader coverage of sequence space, eventually leading to superior generalization if sufficient data are available. Looking ahead, MDC can serve as a foundation for next-generation S2F modeling approaches, including transfer learning or a source to pre-train comprehensive models of transcriptional regulation.

## Methods

### Data pre-processing

For the input, the “assayed” sequence and any specified upstream or downstream scaffolds were concatenated into a single unified sequence in string format, which was converted to an one-hot encoded matrix of shape (L, 4). Missing nucleotides and other syntax designated for multiple nucleotides were represented as equal probabilities for relevant nucleotides. When native sequences were provided as loci on the reference genome, we retrieved the corresponding reference genome and obtained the test sequence based on its index. For datasets consisting of sequences of uneven lengths, shorter sequences were symmetrically padded to match the longest ones.

For the output, the raw data were pre-processed according to the type of the MPRA experiment. For datasets with FACS as the readout assay, the original bin indices were retained as output values and used as indicators of transcriptional efficiency^4,5,18^. For datasets with RNA-seq as the readout assay, a log transformation was consistently applied, specifically computing log_2(#RNA/#DNA), which was then used as the measure of transcriptional efficiency. In cases where the dataset measured alternative splicing efficiency, the ratio of alternative splicing events within the specified regions was computed, with this value, ranging from 0 to 1, used as the output^3,12,13^.

### DNA foundation model embeddings

For each selected dataset in **Table 2**, 256 sequences were sampled as the representative subset. HyenaDNA^19^ was employed to compute the sequence embeddings in the latent space. These DNA foundation models took one-hot encoded sequences as the input and generated high dimensional 1D sequence embeddings as the output. The 1D sequence embeddings were aggregated using mean pooling and were then visualized on the 2D plane using dimension reduction techniques such as t-SNE^22^ (**Figure 1a**).

The specific checkpoints used for DNA foundation model: HyenaDNA: LongSafari/hyenadna-large-1m-seqlen-hf

### Scaling law curve experiments

Based on initial cross-validation experiments, we use a “medium-depth” residual convolutional neural network for S2F models, consisting of convolutional layers with residual connections and a 1D pooling layer followed by fully-connected layers as the output head. Adam optimizer and CyclicLR scheduler were employed to control proper learning rates.

Mean Square Error (MSE) was used as the loss function. After every fixed number of steps, we evaluated the current model on the validation set and computed the Pearson Correlation Coefficient (PCC). Training process was terminated once the validation metric stopped improving for a fixed number of evaluations, and the best saved model was evaluated on the test set.

The detailed hyperparameters were specified below:

- Input head convolutional layer kernel size: 13
- Input head convolutional layer padding: 6
- Other convolutional layers kernel size: 3
- Other convolutional layers padding: 6
- Convolutional layers channels: 256
- Convolutional block: BatchNorm1d+Conv+ReLU
- Pooling layer type: max
- Pooling layer window size: 4
- Adam optimizer learning rate: 1e-4
- Adam optimizer weight decay: 1e-5
- CyclicLR scheduler learning rate max: 3e-4
- CyclicLR scheduler learning rate min: 3e-5
- Validation interval: every 10 training steps
- Early stopping criteria: no improving for 30 evaluations

For each dataset collected, we kept a fixed testing set, which was either sampled or selected based on chromosome index to avoid information leakage (for native sequences). Specifically, we selected chromosome index 4, 9, 14, 19 or sampled 20% sequences when no chromosome index was available as the testing set. Similarly, chromosome index 3, 8, 13, 18 or another 20% were used as the validation set. With the rest left as the training set, subset of varied sizes [1, 4, 16, 64, 256, 1024, 4096, 8192, 16384, 32768, 65536, 131072, 262144] were sampled where larger subsets always contained smaller subsets (skipped last ones when no sufficient sequences). The same S2F model was trained on the same subset 5 times with distinct model initializations to reduce variance.

### Model architecture experiments

Datasets Promoter_1, Enhancer_4, SS_3, and UTR_1r were used in model architecture scaling law curves experiments^4,10,13,14^ (**Table 3**).

Depth: we altered the number of total convolutional blocks used in the S2F model, from 4 layers in the original model, to 2 layers in the “shallow” model, while keeping the other hyperparameters the same (**Figure 2b**). We also altered the number to 1 layer or 9 layers (**Figure S1b**).

Width: we altered the number of channels used in each convolutional layer, from 256 in the original model, to 64 in the “narrow” model, while keeping the other hyperparameters the same (**Figure 2b**). We also altered the number to 128 or 384 (**Figure S1c**).

Transformers: All but the first convolutional blocks used in the original model were replaced by the transformer blocks together with an extra positional encoding. We employed the implementation torch.nn.TransformerEncoderLayer by PyTorch, and assigned d_model as 256, nhead as 4, dim_feedforward as 1024, to ensure its parameter number was comparable with the original convolutional block (**Figure 2c**).

### Cross-species generalization testing

Random vs Native: Both synthetic and natural yeast 5’ UTR sequences were used ^12^. The random dataset UTR_1n consists of 489,242 random sequences of length 50 and the native dataset UTR_1r consists of 11,962 yeast 5’ UTR of length from 2 to 50. No chromosome index was provided for both datasets, so 20% sequences were sampled and used as the same testing set for two scaling law curves drawn (**Figure 3a, Table 4**).

Yeast vs Human: Besides the datasets used above, a human 5’ UTR dataset was used^15^. Specifically, we employed the dataset UTR_2, which consists of 351,575 sequences (a combination of random and native sequences). Similarly, 20% sequences were sampled as the same testing set for all scaling law curves drawn. In addition to those two dataset discussed above, we also combined them together to form two more datasets, which are: (1) a “mixed” dataset UTR_1m, that consists all 501,204 sequences from both datasets, and, (2) a “balanced” dataset UTR_1b, a more balanced combination, which consists of all 11,962 sequences from native dataset and 47,848 sequences sampled from random dataset (**Figure 3b, Table 4**).

## Data and Code availability

The MPRA Dataset Collection and scripts to generate all the figures are available at https://github.com/mostafavilabuw/MPRACollection.

---

**Figure S1.**
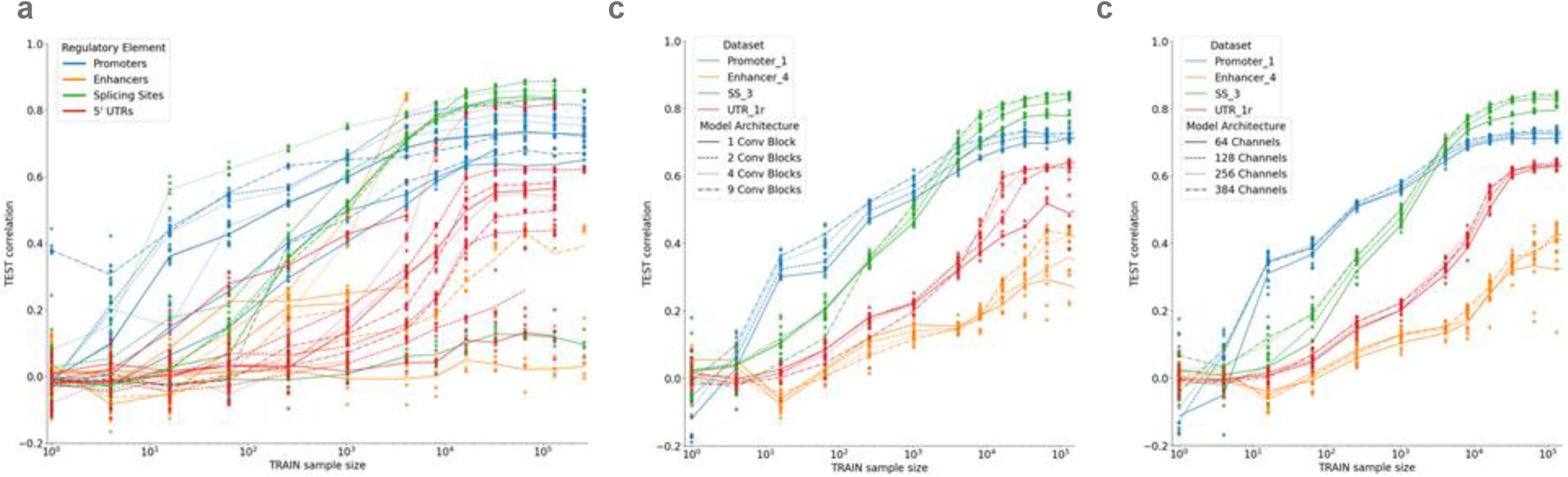
a) Scaling law experiments for almost all datasets in MDC. b) Different line styles represent variations in the number of convolutional blocks. c) Different line styles represent variations in the number of channels/kernels in the first convolutional layer. See **Table 3** for datasets showcased here.

